# Effect of *PPARGC1A* Gly428Ser (rs8192678) polymorphism on athletic sports performance: A meta-analysis

**DOI:** 10.1101/365270

**Authors:** Phuntila Tharabenjasin, Noel Pabalan, Hamdi Jarjanazi

## Abstract

**Background:** Genetics plays a role in sports performance (SP) which, in this study, is measured by power and endurance activities as well as a mix between the two. However, variable results from genetic association studies warrant a meta-analysis to obtain more precise estimates of the association between *PPARGC1A* Gly482Ser polymorphism and SP.

**Methods:** Multi-database literature search yielded 14 articles (16 studies) for inclusion. Pooled odds ratios (ORs) and 95% confidence intervals (CI) were used in estimating associations. Summary effects were modified based on statistical power. Subgroups were SP (power, endurance and mixed) and race (Caucasians and Asians). Heterogeneity was assessed with the I^2^ metric and its sources examined with outlier analysis.

**Results:** Gly allele effects significantly favoring SP (OR > 1.0, P < 0.05) form the core of our findings in: (i) the homogeneous overall effect at the post-modified, post-outlier level (OR 1.13, 95% CI 1.03-1.25, P = 0.01, I^2^ = 0%); (ii) initially homogeneous power SP (ORs 1.22-1.25, 95% CI 1.05-1.44, P = 0.003-0.008, I^2^ = 0%) which precluded outlier treatment; (iii) pre-outlier Caucasian outcomes (ORs 1.29-1.32, 95% CI 1.12-1.54, P = 0.0005) over that of Asians with a pooled null effect (OR 0.99, 95% CI 0.72-1.99, P = 0.53-0.92); (iv) homogeneous all > 80% (ORs 1.19-1.38, 95% CI 1.05-1.66, P = 0.0007-0.007, I^2^ = 0%) on account of high statistical power (both study-specific and combined). In contrast, none of the Ser allele effects significantly favored SP and no Ser-Gly genotype outcome favored SP. The core significant outcomes were robust and comparisons subjected to publication bias test showed no evidence of it.

**Conclusion:** Lines of evidence show that the Gly allele effects favor SP. These were observed in the overall, Caucasians and statistically powered comparisons which exhibited consistent significance, stability, robustness, precision and lack of bias, all of which provide good evidence of associations between the Gly allele and SP.

**List of abbreviations:**

A: Asian
ACE: Angiotensin converting enzyme
ACTN3: Alpha (α)-actinin 3
AM: Analysis model
ANOVA: Analysis of variance
df: Degree of freedom
ES: Elevated significance
C: Caucasian
CB: Clark-Baudouin
CI: Confidence interval
CID: Confidence interval difference
E: Endurance
EH: Eliminated heterogeneity
F: Fixed-effects
Fs: Favoring SP
Ds: Disfavoring SP
Gly: Glycine
GS: Gain in significance
Het: Heterogeneity
HWE: Hardy-Weinberg Equilibrium
I^2^: Measure of variability expressed in %
K: Number designation of the study
Li: Lithuania
Log OR: Logarithm of OR
M: Mixed
maf: Minor allele frequency
M-H: Mantel-Haenszel
N: Null
NA: Not applicable
n: Number of studies
OR: Odds ratio
P: P-value
P^a^: P-value for association
P^b^: P-value for heterogeneity
P_o_: Poland
*PPARGC1A*: *Peroxisome proliferator-activated receptor gamma co-activator-1-alpha*
PRISMA: Preferred Reporting Items for Systematic Reviews and Meta-Analyses
PRO: Pre-outlier
PSO: Post-outlier
Pw: Power
R: Random-effects
[R]: Reference number
RH: Reduced heterogeneity
RNS: Retained non-significance
RS: Retained significance
Ru: Russia
SD: Standard deviation
SE: Standard error
Ser: Serine
Sig: Significance
SP: Sports performance

## Introduction

Sports performance (SP) has a highly polygenic and complex phenotype as well as having a multifactorial etiology where genetic and environmental factors contribute to differences among trained athletes [1]. In this study, the role of genetics in SP is examined in the context of strength/power and endurance activities as well as SP activities between the two (mix). The mix represents a continuum of SP activities between power and endurance. Explosive activities such as sprint and weightlifting characterize strength/power while endurance involves slow-burn activities such as marathons and cycling. Power phenotypes comprise characteristics of muscle strength and sprint performance while endurance phenotypes include maximal oxygen uptake and economy of movement [2]. Muscle fiber composition in these two types of athletes differs where activation of types II and I fibers occur during high-intensity activity in power and endurance performances, respectively [3, 4]. These contrasts stem from diverging genetic backgrounds of power and endurance athletes that drive their physiology into different trajectories [5].

Multiple genetic variants are thought to influence muscle function and SP phenotypes [2]. Among the genetic loci associated with SP, *peroxisome proliferator*-*activated receptor gamma co*-*activator*-*1*-*alpha* (*PPARGC1A* or PGC-1α) aroused interest for the varied functions of the proteins it encodes. *PPARGC1A* is encoded by the gene *PPARGC1A* in humans which is crucial in training-induced muscle adaptation because it co-activates a broad range of transcriptional factors that control myriad biological responses [6]. Studies have shown that several amino acid polymorphic sites exist within the coding region of *PPARGC1A*, including Gly482Ser (rs8192678), which is reported to have functional relevance [7].

Current understanding of Gly482Ser in *PPARGC1A* is viewed in terms of its impact on health (e.g. diabetes and obesity) and on athletic phenotype (e.g. endurance sports). In terms of health impact, physiological evidence has shown that Gly482Ser affects blood lipid levels and insulin sensitivity. Compared with carriers of 482Gly, those with 482Ser have higher levels of low density lipoprotein cholesterol [8] and higher insulin resistance [9]. As a result, such persons have increased risk for Type 2 diabetes [10, 11]. A number of similar and variable findings on this topic have been reported in the literature that warranted coverage in published meta-analyses [12-14]. Research at the meta-analysis level indicates sufficient number of Gly482Ser studies to warrant this treatment.

It shows the prevailing knowledge on this polymorphism tending more toward health rather than SP. Gly482Ser studies in SP have been heterogeneous given the various research approaches, variable sample sizes and different population profiles that characterize them. Such heterogeneity then renders the proposed associations to be inconsistently replicated. Clearly, utilizing research methods that synthesize diverging primary study results is needed which meta-analysis seems most suitable to resolve. To the best of our knowledge, this is the first meta-analysis to examine the role of Gly482Ser in SP. In this study, we focus on the genetic role of this polymorphism in SP by using an array of meta-analytical techniques such as a scale to evaluate quality of the primary literature, tests of association, outlier and modifier treatments, sensitivity analysis and tests for publication bias in order to assess the strength of evidence. This in-depth treatment precludes covering other SP related genes. Thus, we view the single-polymorphism approach most suitable for reasons of brevity and clarity of reporting.

## Materials and methods

### Selection of studies

We searched MEDLINE using PubMed, Science Direct and Google Scholar for association studies as of June 15, 2018. Terms used were “*peroxisome proliferator*-*activated receptor*-*gamma co*-*activator 1*-*alpha*”, “*PPARGC1A*”, “PGC1-α”, “rs8192678”, “sports performance”, and “polymorphism” as medical subject heading and text, restricted to the English language.

References cited in the retrieved articles were also screened manually to identify additional eligible studies. Inclusion criteria were: (i) case–control study design evaluating the association between *PPARGC1A* polymorphisms and SP; (ii) studies comply with the Hardy-Weinberg Equilibrium (HWE); (iii) sufficient genotype or allele frequency data to allow calculation of odds ratios (ORs) and 95% confidence intervals (CIs). Exclusion criteria include: (i) studies that do not involve SP (e.g. *PPARGC1A* polymorphism effects in pathophysiological conditions such as diabetes or cases were non-athletes); (ii) studies whose genotype or allele frequencies were otherwise unusable / absent or when available but combined with other polymorphisms, preventing proper data extraction; (iii) in case of duplicates, we chose the most recent article; (iv) reviews; (v) no controls; (vi) when controls were present, their frequencies deviated from the HWE; (vii) non-human subjects and non-English articles.

### Data extraction

Two investigators (PT and NP) independently extracted data and arrived at consensus. The following information was obtained from each publication: first author’s name, publication year, country of origin and SP type. S1Table tabulates information on the quantitative data. The core information here is the genotype data, which was used to conduct the meta-analysis. This thus precludes extraneous information such as environmental data which was either unmentioned or unquantified in the primary literature. Of note, HWE was assessed using the application in https://ihg.gsf.de/cgi-bin/hw/hwa1.pl. Authors were contacted in order to obtain more information on incomplete data. Responses to our requests have not been received thus far.

**<u>S1 Table</u> Quantitative characteristics of *PPARGC1A* studies examining associations with sports performance**

### Quality assessment of the studies

We used the Clark-Baudouin (CB) scale to evaluate methodological quality of the included studies [15]. Suitability of this scale is based on criteria such as P-values, statistical power, corrections for multiplicity, comparative sample sizes between cases and controls, genotyping methods and HWE, features found in the component articles. CB scores range from zero (worst) to 10 (best) where scoring is based on quality (low: < 5, moderate: 5-7, and high: ≥ 8).

### Meta-analysis

Gly482Ser associations with SP (OR) were estimated for each study. Where genotype values were zero, we applied Laplace correction by adding a pseudo-count of one to all values of the data set [16] before generating the forest plots. We used the allele-genotype approach to enable comparison with study-specific outcomes. We thus compared the following for Gly482Ser: (i) Gly allele with Ser-Gly/Ser-Ser genotype; (ii) Ser allele with Ser-Gly/Gly-Gly genotype; (iii) Gly/Ser genotype with homozygous Gly-Gly and Ser-Ser genotypes. To compare effects on the same baseline, we used raw data for genotype frequencies to calculate pooled ORs. Pooled ORs with their accompanying 95% CIs provide indications that assess the strength of evidence. These are: (i) magnitudes of effects are higher or lower when the values are farther from or closer to the OR value of 1.0 (null effect), respectively; (ii) OR values are significant when P < 0.05 (two-sided); (iii) CID (confidence interval difference) results when the lower CI is subtracted from the upper CI, which indicates precision of effects. High and low CID values indicate low and high precision, respectively.

Heterogeneity between studies was estimated with the χ^2^-based Q test [17], explored with subgroup analysis [17] and quantified with the I^2^ statistic which measures degree of inconsistency between studies [18]. The fixed effects model [19] was used when P ≥ 0.10 or I^2^ < 50%, otherwise we opted for the random effects model [20], signifying presence of heterogeneity. We identified sources of heterogeneity using the Galbraith plot [21] to detect outliers, then followed by re-analysis. Three features of this re-analysis are worth noting: (i) outlier treatment dichotomizes the comparisons into pre-outlier (PRO) and post-outlier (PSO) which is visualized with use of the forest and Galbraith plots as well as integrated in the design of the summary tables; (ii) outlier treatment is applied in PRO which assumes the random-effects status; (iii) PSO outcomes are fixed-effects, where larger studies are accorded more weight [22]. The Bonferroni correction, applied to independent comparisons only, was used to adjust for multiple testing. Sensitivity analysis, which involves omitting one study at a time and recalculating the pooled OR, was used to test for robustness of the summary effects. Publication bias was assessed on comparisons with ≥ 10 studies only [23]. Data were analyzed using Review Manager 5.3 (Cochrane Collaboration, Oxford, England), SIGMASTAT 2.03, SIGMAPLOT 11.0 (Systat Software, San Jose, CA) and WINPEPI [24].

## Results

### Search results

Fig 1 outlines the study selection process in a flowchart following PRISMA (Preferred Reporting Items for Systematic Reviews and Meta-Analyses) guidelines. Initial search resulted in 523 citations, followed by a series of omissions (S2 Table) that eventually yielded 14 articles for inclusion [25-38]. Of the 14, two articles [29, 34] presented independent data from two populations placing the total number of studies to 16 (Table 1).

**Fig 1.**
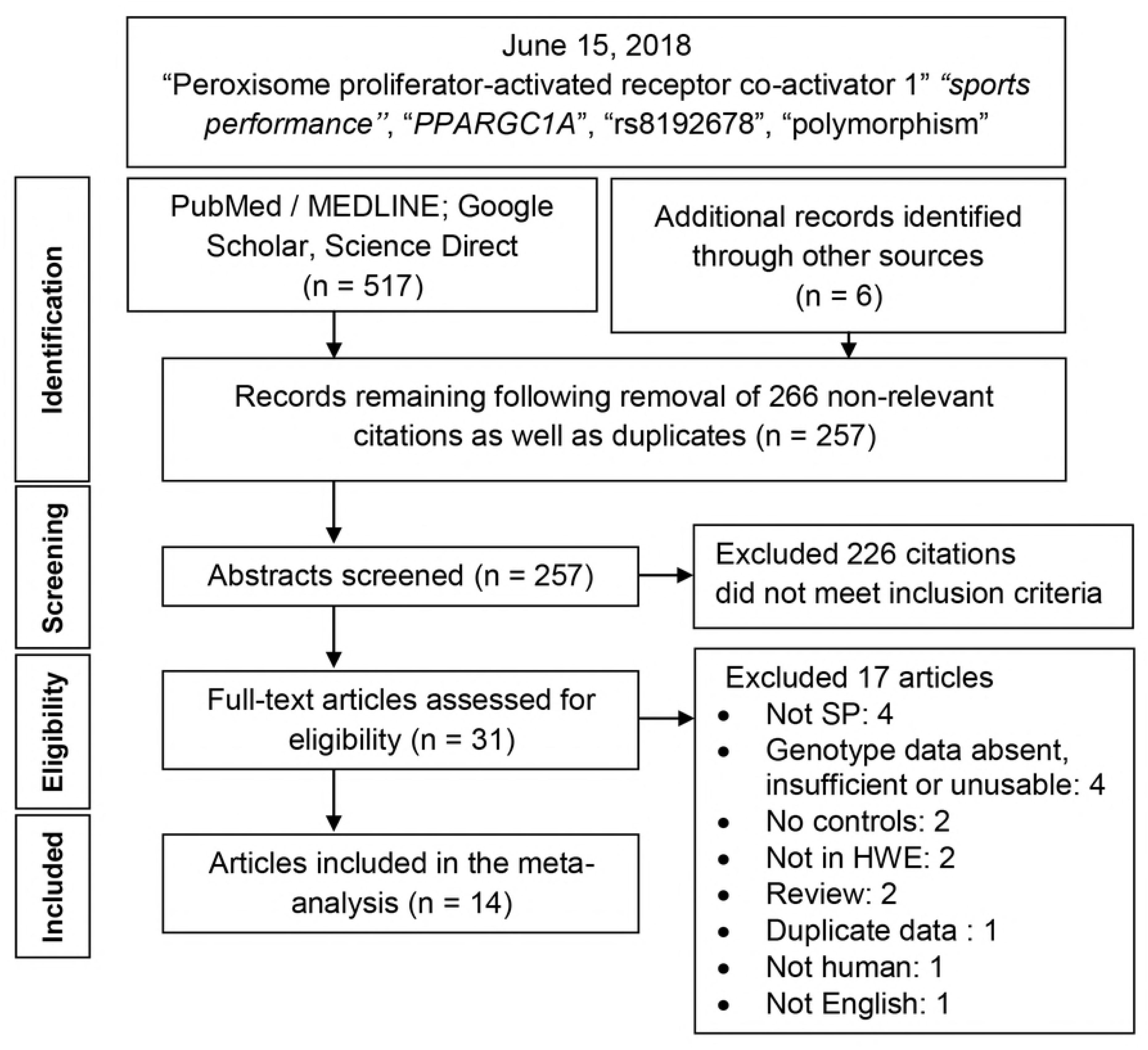
Summary flowchart of literature search.

**Table 1.** Characteristics of the included studies that examined the association of *PPARGC1A* Gly482Ser polymorphism with sports performance.

| K | First author | [R] | Year | n | Country | Race | SP status | CB score |
| --- | --- | --- | --- | --- | --- | --- | --- | --- |
| 1 | Ahmetov | [25] | 2009 | 1 | Russia | C | E | 7 |
| 2 | Eynon | [26] | 2011 | 1 | Israel | C | Pw/E | 8 |
| 3 | Gineviciene 1 | [27] | 2011 | 1 | Lithuania | C | Pw/E/M | 9 |
| 5 | Gineviciene 2 | [28] | 2012 | 1 | Lithuania | C | M | 7 |
| 4 | Gineviciene 6 | [29] | 2016 | 2 | Lithuania/Russia | C | Pw | 8 |
| 6 | Grealy | [30] | 2015 | 1 | Australia | C | E | 7 |
| 7 | He | [31] | 2014 | 1 | China | A | E | 5 |
| 8 | Jin | [32] | 2014 | 1 | Korea | A | M | 7 |
| 9 | Lucia | [33] | 2005 | 1 | Spain | C | E | 7 |
| 10 | Maciejewska | [34] | 2012 | 2 | Poland/Russia | C | Pw/E/M | 7 |
| 11 | Maruszak | [35] | 2012 | 1 | Poland | C | Pw/E | 5 |
| 12 | Muniesa | [36] | 2010 | 1 | Spain | C | E | 7 |
| 13 | Peplonska | [37] | 2016 | 1 | Poland | C | M | 7 |
| 14 | Yvert | [38] | 2016 | 1 | Japan | A | E | 8 |

K: number designation of the article; [R]: reference number; n: number of studies; C: Caucasian; A: Asian; SP: sports performance; E: endurance; M: mixed; Pw: power; CB: Clark-Baudouin

**Fig 1 Summary flowchart of literature search**

<u>S2 Table</u> **List of excluded articles**

### Characteristics of the included studies

Table 1 shows that participants in most of the studies were Euro-Slavic with three (Australia, Japan and Korea) contributing to geographical heterogeneity [30, 32, 38]. Subgroups by sport type and race comprised of power [26, 27, 29, 34, 35], endurance [25-27, 30, 31, 33-36, 38] and mixed [27, 28, 32, 34, 37], Caucasian [25-30, 33-37] and Asian [31, 32, 38], respectively. Median and range CB score of 7.0 (5-9) indicates that methodological quality of the component studies was high. S1 Table shows that the meta-analysis is composed of seven, 11, and six studies in power, endurance, and mixed, respectively. Quantitative features include sample sizes, genotype frequencies in cases/controls and minor allele frequencies (maf) in each of the sport types (S1Table). The maf means and standard deviations of Caucasians (0.34 ± 0.05) and Asians (0.49 ± 0.05) differed significantly (t = −4.86, P < 0.001). The checklists for PRISMA and meta-analysis for genetic association detailed features of this meta-analysis in accordance with the guidelines (S3 and S4 Tables).

<u>S3 Table</u> **PRISMA checklist**

<u>S4 Table</u> **Checklist of meta-analysis for genetic association studies**

### Meta-analysis outcomes

Table 2, S5 and S6 Tables summarize the meta-analysis outcomes by order of genetic comparisons (Gly and Ser alleles and Ser-Gly genotype). Between these three tables, number of pooled ORs > 1.0 (favoring SP) was most in Gly allele and least in Ser allele and none in Ser-Gly genotype. This positions the Gly allele analysis as central to our findings because it presents the most convincing evidence indicating the favoring of SP.

**Table 2.**
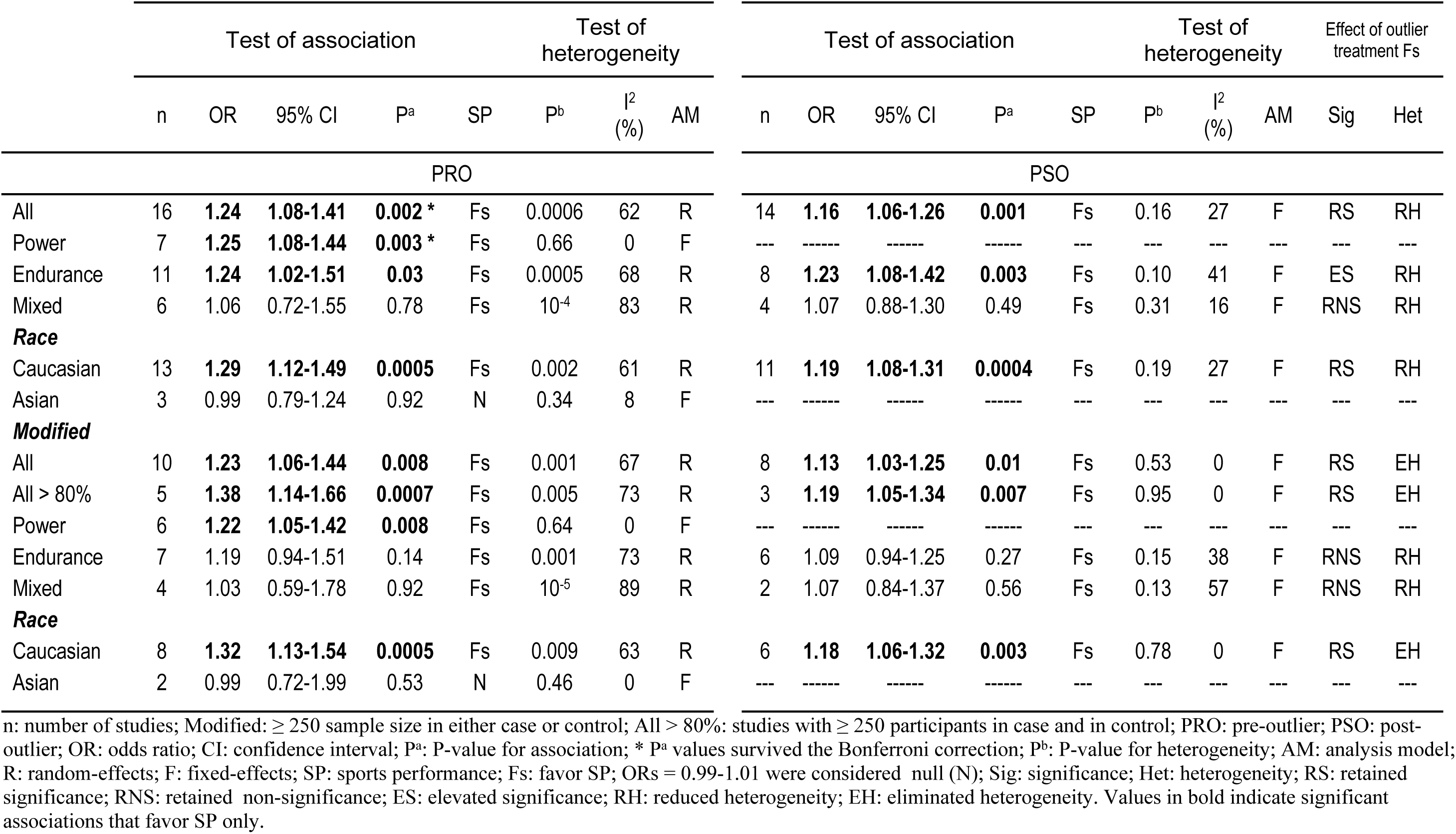
Outlier and modified effects for Gly allele associations with sports performance.

<u>S5 Table</u> **Outlier and modified effects for Ser allele associations with sports performance**

<u>S6 Table</u> **Modified effects for Ser-Gly genotype associations with sports performance**

### Gly allele effects

Table 2 shows the Gly allele associations where 20 (91%) of the 22 comparisons favored SP (OR >1.0). Of the 20 SP favoring outcomes, 14 (70%) were statistically significant (P < 0.05). These SP favoring and significant features were observed in the overall (ORs 1.16-1.24, 95% CI 1.06-1.41, P = 0.001-0.002) and Caucasian subgroup (ORs 1.19-1.29, 95% CI 1.08-1.49, P = 0.0004-0.0005). In contrast to Caucasians, the Asian effects were null and non-significant (OR 0.99, 95% CI 0.79-1.24, P = 0.92). Fig 2 shows the pooled findings in power SP (OR 1.25, 95% CI 1.08-1.44, P = 0.003) which were initially homogeneous (I^2^ = 0%) precluding outlier treatment. PRO effects on endurance (ORs 1.19-1.24, 95% CI 0.94-1.51, P = 0.03-0.14) and mixed (ORs 1.03-1.06, 95% CI 0.59-1.78, P = 0.78-0.92) were more modulated than the power outcomes (Table 2).

**Fig 2.**
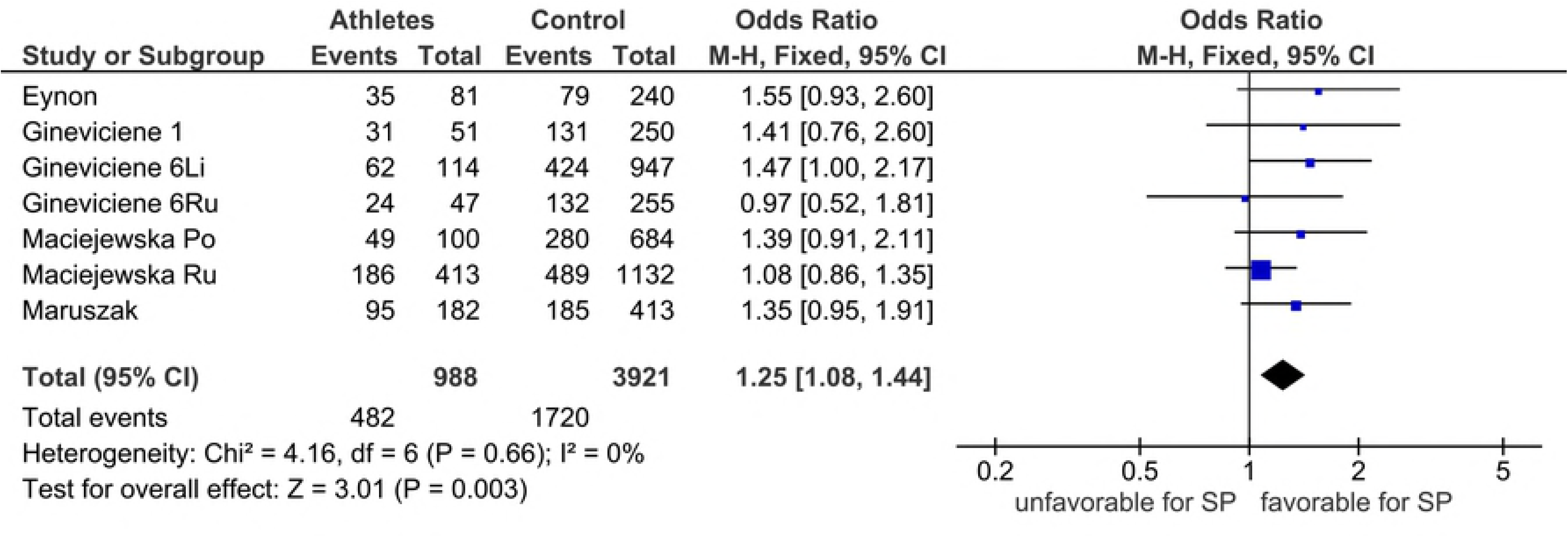
Forest plot outcome of *PPARGC1A* Gly allele effects on SP power in the premodifier analysis. Diamond denotes the pooled odds ratio (OR). Squares indicate the OR in each study, with square sizes directly proportional to the weight contribution (%) of each study. Horizontal lines represent 95% confidence intervals (CI). The Z test for overall effect indicates significance (P = 0.003). The chi-square test shows absence of heterogeneity (P = 0.66, I^2^ = 0%). M-H: Mantel-Haenszel; CI: confidence interval; df: degree of freedom; I^2^: measure of variability expressed in %.

Outlier treatment had multiple effects on a number of parameters: (i) heterogeneity was reduced (P_heterogeneity_ ≥ 0.10) or eliminated (I^2^ = 0%); (ii) significance was retained (overall, Caucasian, all > 80%) and elevated (endurance); (iii) precision of effects was increased (reduction of CID values from PRO to PSO).

The mechanism of outlier treatment is visualized in Figs 3-5. Fig 3 shows the following features in the Gly allele endurance PRO analysis: (i) heterogeneous (P_heterogeneity_ = 0.0005, I^2^ = 68%); (ii) moderately significant (OR 1.24, 95% CI 1.02-1.51, P = 0.03); (iii) CID of 0.49 (CI 1.02-1.51). In Fig 4, the Galbraith plot identifies three studies as the outliers [25, 30, 38] located above the +2 and below the −2 confidence limits. In Fig 5, the PSO outcome (outliers omitted) shows reduced heterogeneity (P_heterogeneity_ = 0.10, I^2^ = 41%), increased significance (OR 1.23, 95% CI 1.08-1.42, P = 0.003) and increased precision with a reduced CID of 0.34 (CI 1.08-1.42). This operation is numerically summarized in Table 2.

**Fig 3.**
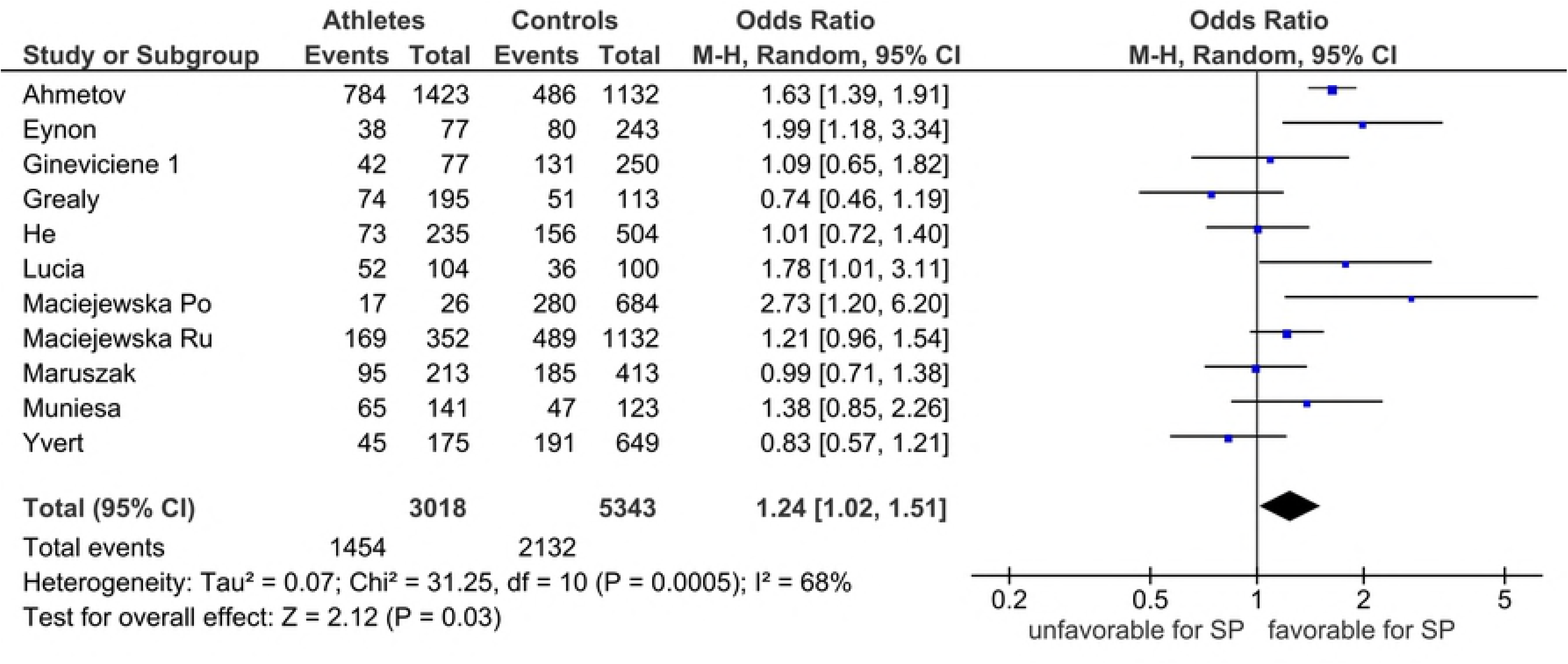
Forest plot outcome of *PPARGC1A* Gly allele effects on SP endurance in the PRO analysis. Diamond denotes the pooled odds ratio (OR). Squares indicate the OR in each study, with square sizes directly proportional to the weight contribution (%) of each study. Horizontal lines represent 95% confidence intervals (CI). The Z test for overall effect indicates significance (P = 0.03). The chi-square test shows presence of heterogeneity (P = 0.0005, I^2^ = 68%). M-H: Mantel-Haenszel; CI: confidence interval; df: degree of freedom; I^2^: measure of variability expressed in %.

**Fig 4.**
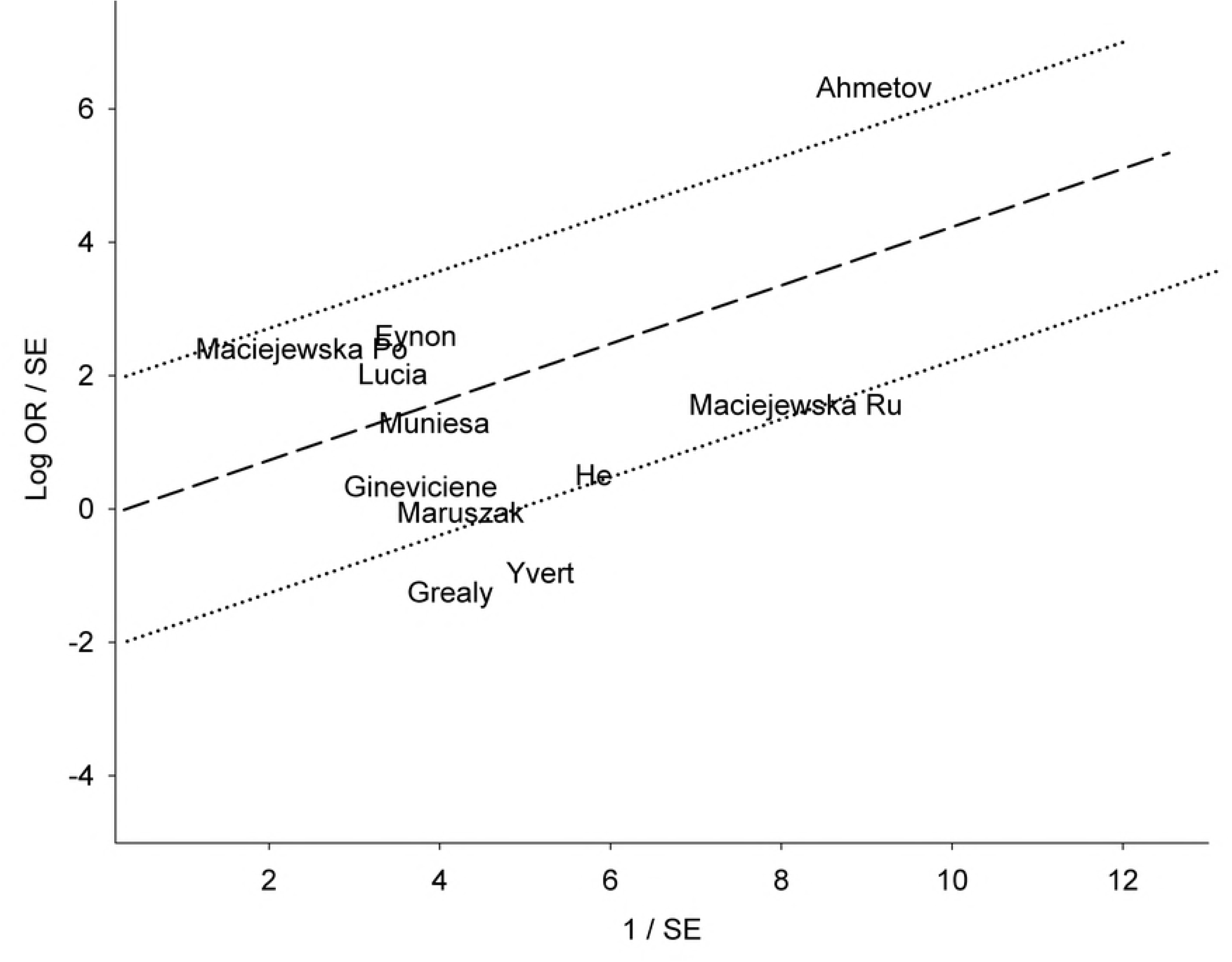
Galbraith plot analysis of Gly allele effects on SP in endurance identifying the sources of heterogeneity. Log OR: logarithm of standardized odds ratio; SE: standard error. The three studies that lie above and below the +2 and - 2 confidence limits are the outliers.

**Fig 5.**
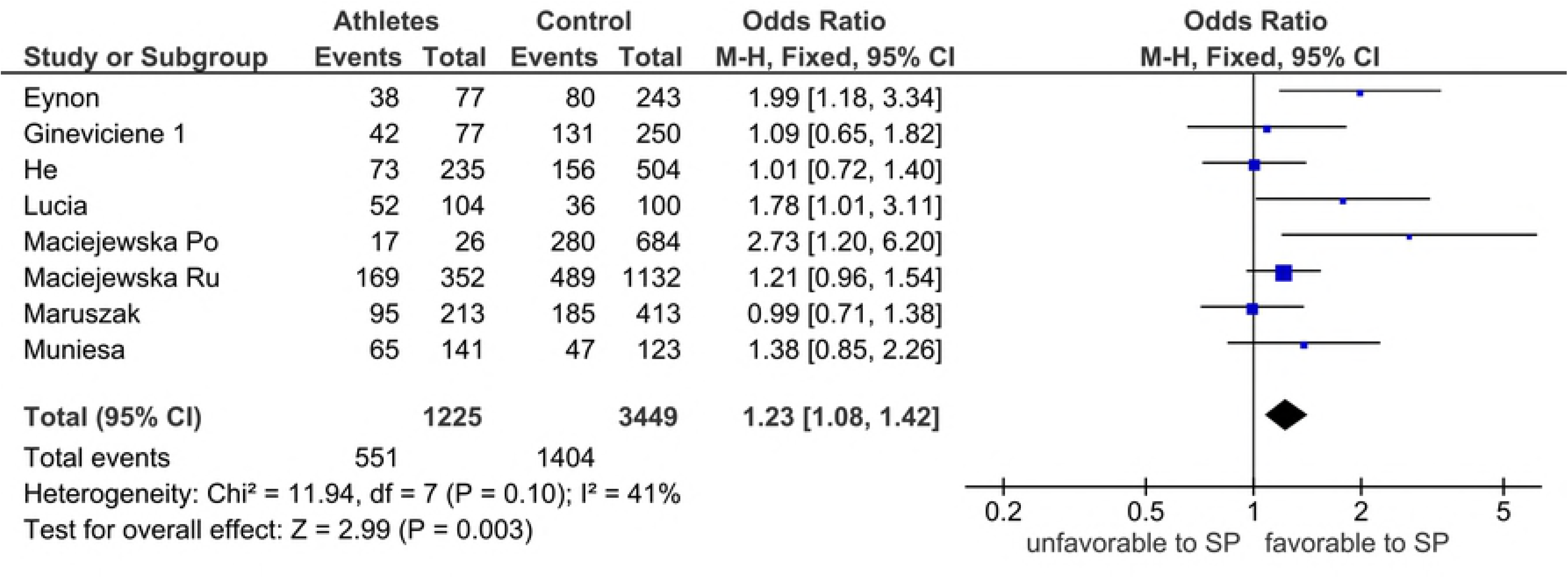
Forest plot outcome of outlier treatment on *PPARGC1A* Gly allele effects on SP endurance in the PSO analysis. Diamond denotes the pooled odds ratio (OR). Squares indicate the OR in each study, with square sizes directly proportional to the weight contribution (%) of each study. Horizontal lines represent 95% confidence intervals (CI). The Z test for overall effect indicates high significance given (P = 0.003). The chi-square test indicates non-heterogeneity (P = 0.10, I^2^ = 41%). M-H: Mantel-Haenszel; CI: confidence interval; df: degree of freedom; I^2^: measure of variability expressed in %.

**<u>Fig 2</u> Forest plot outcome of *PPARGC1A* Gly allele effects on SP power in the premodifier analysis.**

**<u>Fig 3</u> Forest plot outcome of *PPARGC1A* Gly allele effects on SP endurance in the PRO analysis.**

**<u>Fig 4</u> Galbraith plot analysis of Gly allele effects on SP in endurance identifying the sources of heterogeneity.**

**<u>Fig 5</u> Forest plot outcome of outlier treatment on *PPARGC1A* Gly allele effects on SP endurance in the PSO analysis.**

### Modified effects in Gly comparisons

Testing a single nucleotide polymorphism using a case-control design has been calculated to require a sample size of 248 participants in each group to achieve a statistical power of 80% [39]. To approximate this level and still achieve enough studies, we selected those with at least 248 participants in either cases or controls for Gly allele comparison only. However, we also included five studies from four papers [25, 34, 35, 37] with > 248 participants both in cases and in controls which we termed “all > 80%”. Using the G*Power program [40], statistical powers in each of these three studies were calculated to range between 81.0% and 99.9% assuming an α level of 5%. Outcomes of all > 80% (ORs 1.19-1.38, 95% CI 1.05-1.66, P = 0.0007-0.007) were not only homogeneous (I^2^ = 0%) at the PSO level.

Modified analysis generated the following outcomes for other comparisons (PRO and PSO) with elimination of both heterogeneity (Overall, Caucasian and Asia: I^2^ = 8-27% to 0%) and significance (P = 0.03 to 0.14) in endurance SP. However, significance was retained in overall, power SP and Caucasians (Table 2).

### Ser allele and Ser-Gly genotype effects

S5 Table shows the Ser allele associations where three (14%) of the 22 comparisons (PRO and PSO) favored SP (ORs 1.08-1.13, 95% CI 0.83-1.44). Of the three, none were significant (P > 0.05). Eighteen of the 22 (82%) comparisons in PRO and PSO disfavored SP (ORs 0.57-0.95, 95% CI 0.41-1.23) of which, two (11%) were significant (P < 0.05). The remaining comparison was the null Asian effect (OR 1.01, 95% CI 0.79-1.31, P = 0.91).

S6 Table shows the Ser-Gly genotype associations 13 outcomes of which 11 (85%) disfavored SP (ORs 0.80-0.87, 95% CI 0.69-1.00, P < 10^−4^-0.06) and two (15%) had null (Asian) outcomes (ORs 1.00-1.01, 95% CI 0.80-1.26, P = 0.95-1.00). All Ser-Gly genotype comparisons had the fixed-effects feature indicating initial non-heterogeneity.

### Sensitivity analysis and publication bias

Sensitivity analysis was performed using a modified protocol that confined this treatment to the significant Gly allele findings. Pooled effects that retained (P < 0.05) or lost (P > 0.05) significance were considered robust and not robust, respectively. Table 3 identifies the robust and non-robust comparisons. The most robust comparisons were overall and all > 80% in the PRO analysis and all Caucasian outcomes. The PRO analysis had seven robust outcomes and as many interfering studies; PSO had half the number of robust outcomes and one unduplicated interfering study [34]. All unstable comparisons are attributed to six studies from five articles [25, 26, 33, 34, 36].

**Table 3.** Sensitivity analysis of Gly allele comparisons with significant outcomes favoring sports performance.

| Comparison | PRO | PSO | A | B |
| --- | --- | --- | --- | --- |
| Overall | Robust | Robust | 0 | 2 |
| Power | Robust | ---- | 0 | 1 |
| Endurance | [25, 26, 33, 34, 36] | Robust | 5 | 1 |
| Caucasian | Robust | Robust | 0 | 2 |
| **Modified** |  |  |  |  |
| Overall | Robust | [34] | 1 | 1 |
| All $> 80\%$ | Robust | [34] | 1 | 1 |
| Power | Robust | ---- | 0 | 1 |
| Caucasian | Robust | Robust | 0 | 2 |
| A | 5 | 2 |  |  |
| B | 7 | 4 |  |  |

PRO: pre-outlier; PSO: post-outlier; A: number of references that contributed to non-robustness; B: number of robust outcomes

Data (study-specific ORs) used to test for publication bias was determined to be normally distributed (Kolmogorov-Smirnov test: P > 0.05). Hence, we used the Egger’s regression asymmetry test only which showed no evidence of publication bias (Table 4).

**Table 4.**
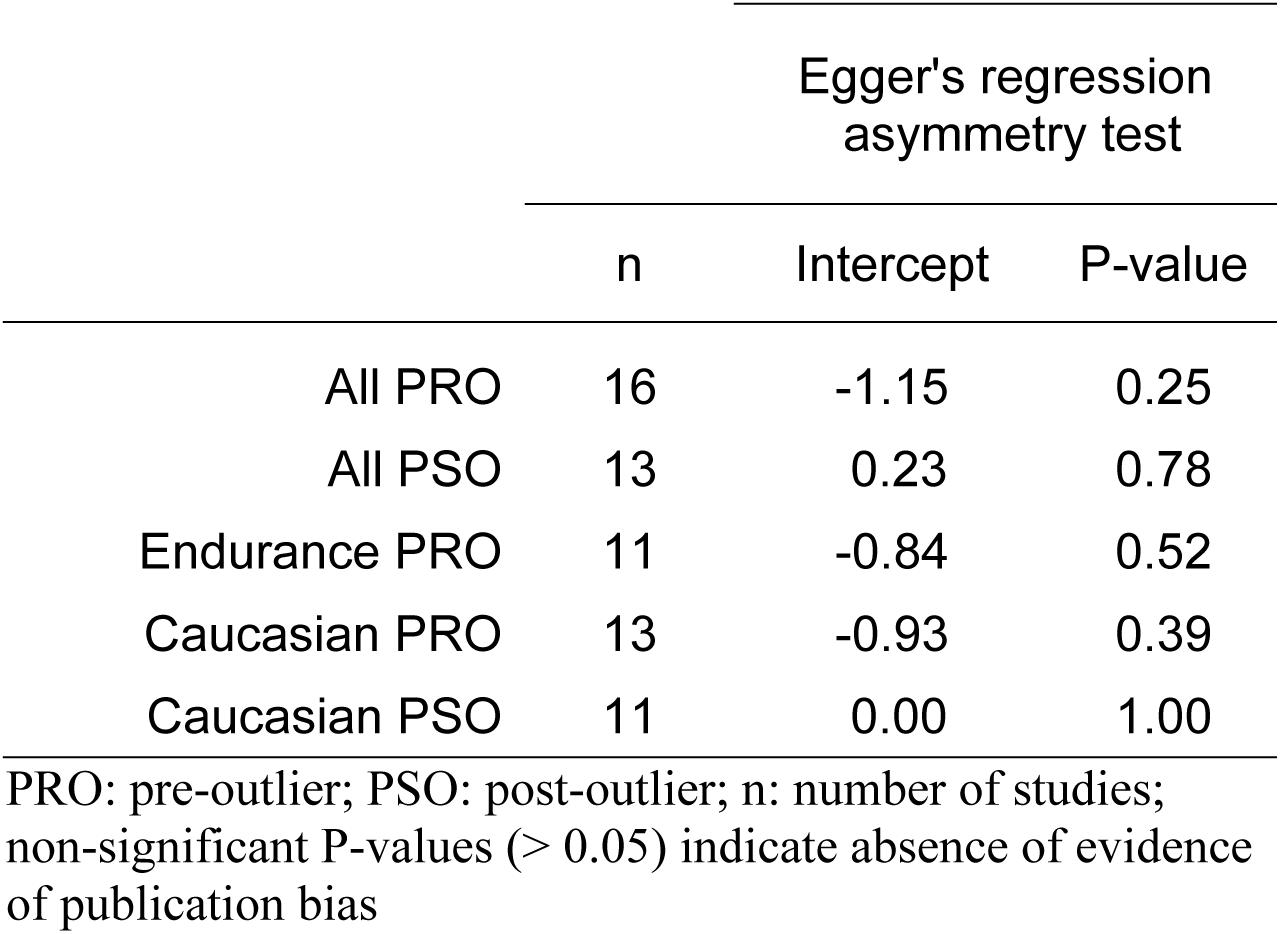
Publication bias assessment of Gly allele comparisons with significant outcomes favoring sports performance.

|  | Egger's regression asymmetry test |  |  |
| --- | --- | --- | --- |
|  | n | Intercept | P-value |
| All PRO | 16 | -1.15 | 0.25 |
| All PSO | 13 | 0.23 | 0.78 |
| Endurance PRO | 11 | -0.84 | 0.52 |
| Caucasian PRO | 13 | -0.93 | 0.39 |
| Caucasian PSO | 11 | 0.00 | 1.00 |

PRO: pre-outlier; PSO: post-outlier; n: number of studies; non-significant P-values ( $> 0.05$ ) indicate absence of evidence of publication bias

## Discussion

We should point out that interpreting effects of polymorphisms differ between disease and SP, besides their respective domains in pathology and in normal phenotype. Disease effects are viewed in terms of protection (reduced risk) or susceptibility (increased risk) both of which have equal importance especially when significant. SP effects on the other hand, are better contextualized when interpreting outcomes that favor SP (OR > 1.0). Thus, our reason for de-emphasizing ORs < 1.0 is that these values disfavoring SP do not contribute to promoting SP.

### Summary of effects

This meta-analysis delineated which genetic component of Gly428Ser in the *PPARGC1A* gene favors SP (Gly allele) and those that do not (Ser allele and Ser-Gly genotype). Subjecting these components to meta-analysis treatments (outlier, modified, and sensitivity) impacted on the outputs. For example, the combined application of outlier and modifier treatments unraveled favorable features of the pooled outcomes that included reduced/ eliminated heterogeneity and elevated statistical power. Outlier treatment attempts to resolve heterogeneity issues that are inherent in meta-analysis. Modifier treatment operates through exclusion of underpowered studies. Underpowered outcomes appear to be common in candidate gene studies [22] and are prone to the risk of Type 1 error. This risk was addressed by generating a comparison with increased statistical power and correcting for multiple comparisons. Thus, the all > 80% modified analysis was created especially in light of significant results [41] and the Bonferroni correction to minimize the possibility of false-positive outcomes [42]. Both outlier and modifier treatments raise the levels of evidence presented here and highlight the transparency of our findings. The main findings of this study center on the Gly allele on account of statistical significance in power SP, all > 80% and the Caucasian subgroup. The SP favoring Gly allele bearing Caucasians but not Asians may be attributed to the significant difference in maf between the two races. While the Asian subgroup acquired zero heterogeneity on account of modifier treatment, the Caucasian subgroup acquired homogeneity on account of modifier and outlier treatments combined.

While homogeneous outcomes in meta-analysis improve the quality of evidence, heterogeneous results are unavoidable and must be addressed. A pro-active approach to addressing heterogeneity is identifying its sources using outlier treatment to re-analyze the results. Our application of outlier treatment had far-reaching effects, impacting on significance, heterogeneity and precision. However, it did not eliminate heterogeneity for the most part. Reduced heterogeneity, notwithstanding, our meta-analysis findings, such as those in the endurance outcomes agree with the physiological evidence [32, 34]. Variable pooled outcomes (ORs that skirt the null effect [ORs 0.99-1.01] and none that indicate favoring SP) observed in mixed sports effectively differs from the SP favoring power outcomes which seem to reflect the inherent phenotypic heterogeneity of this sport type [43]. Ser allele and Ser-Gly genotype effects were consistent in disfavoring SP, regardless of sport type and outlier treatment [34]. Favoring SP outcomes are underpinned by a number of important features: (i) similar repeated effects in the comparisons (consistency); (ii) reduced PSO heterogeneity (outcomes of outlier treatment); (iii) enhanced PSO significance (endurance); (iv) increased precision (reduced CID values from PRO to PSO); (v) robustness (resistance to sensitivity treatment) all of which present strong evidence of Gly428Ser associations with SP.

The *PPARGC1A* gene presents an interesting subject regarding associations between genotypes and SP. In particular, Gly482Ser has been regarded a true if not a promising genetic variant in determining SP status for both power and endurance-type athletes [2]. We performed this meta-analysis because reported findings on the role of Gly428Ser in SP have been variable and the number of articles was ripe for synthesizing the variable findings. Methodological problems may explain the discrepancies, which include limited statistical power, unrecognized confounding factors, misleading definition of phenotypes and stratification of populations [15]. However, variability of the primary study outcomes for Gly428Ser in SP hedge on the type of sport they refer to. For instance, the Ser allele has been found to be less frequent in elite athletes of the endurance [33] and power phenotypes [27]. The Ser allele has been reported to disfavor endurance activities; in contrast to the Gly allele which was found useful [32, 34]. Other reports, however, indicate the Ser allele as useful in power activities [29].

### Genetic and physiological correlates

Physiological stress or increased energy demands such as that elicited by exercise training readily alter expression levels of *PPARGC1A* [44]. *PPARGC1A* is implicated in promoting gene expression and muscle morphology characteristic of type I oxidative fibers in skeletal muscle. Investigators have examined the role of *PPARGC1A* mRNA expression in SP where its levels were impacted by exercise training in both mouse and human skeletal muscle [43]. In mouse muscle, *PPARGC1A* is required to uphold mitochondrial protein expression which in turn is needed for oxidative phosphorylation and perturbation of this cascade results in diminished exercise capacity [45]. In humans, Mathai et al. demonstrated that one session of protracted endurance activity induces elevated transcription and mRNA levels of *PPARGC1A* [46]. *PPARGC1A* is expressed at high levels in metabolically active tissues where mitochondria are abundant and oxidative metabolism is active in brown adipose tissue, the heart, and skeletal muscle, whereas the expression level is low in white adipose tissue, liver, and pancreas [47, 48]. *PPARGC1A* plays a role in the regulation of cellular energy metabolism. Not only has been identified as master regulator of mitochondrial biogenesis, but it has also been shown to regulate proteins involved in angiogenesis and anti-oxidant defense as well as affect expression of inflammatory markers [34, 49]. It mediates skeletal muscle fiber type switching, upregulation of enzymes involved in lipid utilization and β-oxidation, and promotes glucose metabolism through upregulating hepatic gluconeogenic genes [46, 48]. Transition from glycolytic type IIb to mitochondria-rich types IIa and I characterize SP among power athletes. Combination of both peak force/power and ability to sustain high-intensity efforts for extended periods during a competition [4] is the process that uses oxidative metabolism [33, 34]. A main deciding factor of maximal sustainable power is mitochondrial amount in the recruited muscle fibers [50].

### Strengths and limitations

Interpreting our findings here is best done in the context of its limitations and strengths. Limitations of our study include: (i) dominating presence of Slavic Caucasian participants (Russia, Lithuania). This precludes extrapolation of the findings to other ethnic groups. More studies are warranted to better represent a wider range of ethnic subgroups, particularly Asian populations; (ii) we did not examine female effects because of data unavailability. Only one [26] of the component studies presented gender-discriminating data which was insufficient to perform subgroup analysis. Although gender differences are not always clear, genes seem to play a more prominent role in male than in female strength determination [51]; (iii) most of the component studies were underpowered; (iv) heterogeneity of the PRO findings; (v) elevated statistical power through modified treatment was countered by non-robustness in the PSO analysis of overall and all > 80%; (vi) caution maybe warranted in concluding strong associations of the Gly allele in our study, given the possibility that this SP increasing allele may be in linkage disequilibrium with the true functional allele [52].

On the other hand, the following strengths not only add to the epidemiological, clinical and statistical homogeneity (hence, combinability) of the studies, but also minimize bias and underpin the magnitude of associations: (i) the combined sample sizes of the overall and SP types yielded high statistical power (S1 Table); (ii) screening for studies whose controls deviated from HWE effectively corrected for genotyping errors, which minimizes methodological weaknesses [53]; (iii) overall methodological quality (determined by CB) of the included studies was high; (iv) outlier treatment reduce and eliminated heterogeneity. Impact of this treatment on significance and precision is viewed to favor our findings; (v) the PRO overall and power outcomes withstood the Bonferroni correction minimizing the possibility of a Type 1 error; (vi) sensitivity treatment conferred robustness to all overall and Caucasian findings, as well as modified overall and all > 80% in PRO; (vii) absence of evidence of publication bias nullifies the notion that it inflates significant pooled outcomes against non-significant results [22].

## Conclusions

Highlights of our findings rests on the fact that most studies in this study lacked statistical power, but when the data are combined using meta-analysis, clear Gly allele effects are uncovered. Here, we present evidence: (i) where Gly allele outcomes favors SP over that of the Ser allele and not at all for Ser-Gly genotype; (ii) within the Gly allele, Caucasians are affected but not Asians; (iii) Gly allele favors power SP more than that of endurance and not in mixed sports at all. Of note, strength of the power SP in PRO lies in its homogeneity and stability of surviving the Bonferroni correction. (iv) Strength of the Gly allele effect is shown by the all > 80% outcomes showcasing the statistical power of this modified comparison.

We recognize the complexity of SP which involves interactions between genetic and non-genetic factors opening the possibility of environmental involvement in modifying Gly482Ser effects. Gene-gene and gene-environment interactions have been reported to have roles in associations of *PPARGC1A* polymorphisms with SP [5, 29]. While all but one [26] of the 11 articles mentioned gene-environment interaction, only two addressed haplotype analysis [25, 34]. Nevertheless, all but two [33, 34] analyzed polymorphisms in other genes, the most common being *ACE* (angiotensin converting enzyme) and *ACTN3* (α-actinin) in six [25, 27-30, 36] and four articles [28-30, 36], respectively. Including other SP-related genes in our meta-analysis would have been logistically problematic. Additional well-designed studies (including meta-analyses) exploring other parameters would confirm or modify our results in this study and add to the extant knowledge about the association of *PPARGC1A* polymorphism and SP.

**Data Availability Statement:** All relevant data are within the paper and its Supporting Information files.

**Competing Interests:** The authors have declared that no competing interests exist.

**Funding:** This research did not receive any specific grant from funding agencies in the public, commercial, or not-for-profit sectors.

## Supporting Information

S1 Supplementary Table quantitative features (DOCX)

S2 Supplementary List of excluded articles (DOCX)

S3 Supplementary Table PRISMA checklist (DOCX)

S4 Supplementary Table checklist meta-analysis on genetic associations (DOCX)

S5 Supplementary Table Ser allele summary outcomes (DOCX)

S6 Supplementary Table Gly-Ser genotype summary outcomes (DOCX)

## Acknowledgements

We thank Prof. Chumpol Pholpramool for reviewing the final draft

## Author Contributions

**Conceptualization:** NP, PT

**Data curation:** NP, HJ

**Formal analysis:** NP, PT

**Investigation:** PT, NP

**Methodology:** NP, PT, HJ

**Project administration:** NP, HJ

**Resources:** PT, NP, HJ

**Software:** NP, HJ

**Supervision:** NP

**Validation:** PT, NP, HJ

**Visualization:** NP, PT, HJ

**Writing original draft:** NP, PT, HJ

